# Forgetting in *Drosophila* consists of an increase in uncertainty rather than a stochastic loss of memory

**DOI:** 10.1101/2025.06.26.661725

**Authors:** Junjiro Horiuchi, Nozomi Uemura, Shiro Horiuchi, Minoru Saitoe

## Abstract

While forgetting has been studied extensively in various organisms, its precise nature has often been unclear. Here, we used behavioral experiments in *Drosophila* to determine that a significant aspect of forgetting consists of a decrease in the ability of a memory to induce an appropriate behavior. We tested flies for memory retention at various times after training and then separately retested both flies that chose correctly and those that chose incorrectly. Although the ability to choose correctly decreased over time, we could not measure any differences in memory between flies that initially chose correctly and those that chose incorrectly upon retest. This suggests that forgetting is unlikely to consist of a spontaneous loss of a memory but instead consists of a decrease in the probability of flies that remember choosing the correct behavioral response. Thus, although flies maintain memory over time, there is an increase in uncertainty associated with this memory. We find that forgetting of long-term memories and accelerated forgetting in old flies occur in a similar manner.

## Introduction

There are various types of forgetting. Anecdotally, in humans, one type may consist of a complete lack of access to a memory where we may completely forget to buy milk on the way home from work, and only later recall this memory when we arrive home and have nothing to drink. In contrast, a different type of forgetting consists of an increase in the amount of uncertainty associated with a particular memory. In this situation, we can access the memory, but the contents of the memory are unreliable, leading to uncertain memory recall. Thus, we may remember reading a book, but certain aspects of the plot or storyline may become hazy over time. While forgetting has been extensively studied in various animal models, the specific aspects of memory that decrease upon forgetting have not been well characterized.

*Drosophila* have been a highly useful organism for the study of learning and memory, and more recently, forgetting (Berry & Davis, 2014; Gao et al., 2019; Horiuchi, 2019; Quinn, Harris, & Benzer, 1974; Tully & Quinn, 1985). Similar to other animals, *Drosophila* can learn and form memories, which gradually decay over time. This memory decay can be plotted as a memory retention curve or forgetting curve (see Fig. 1) (Tully & Quinn, 1985), and similar to other organisms, as *Drosophila* age, they suffer an accelerated loss of memory (Mery, 2007; Tamura et al., 2003). Thus, forgetting can be reliably measured in *Drosophila*. However, specific characteristics of forgetting in flies, whether it consists of a loss of memory of an association, or whether it consists of an increase in uncertainty regarding an association, have not yet been analyzed.

**Figure 1.**
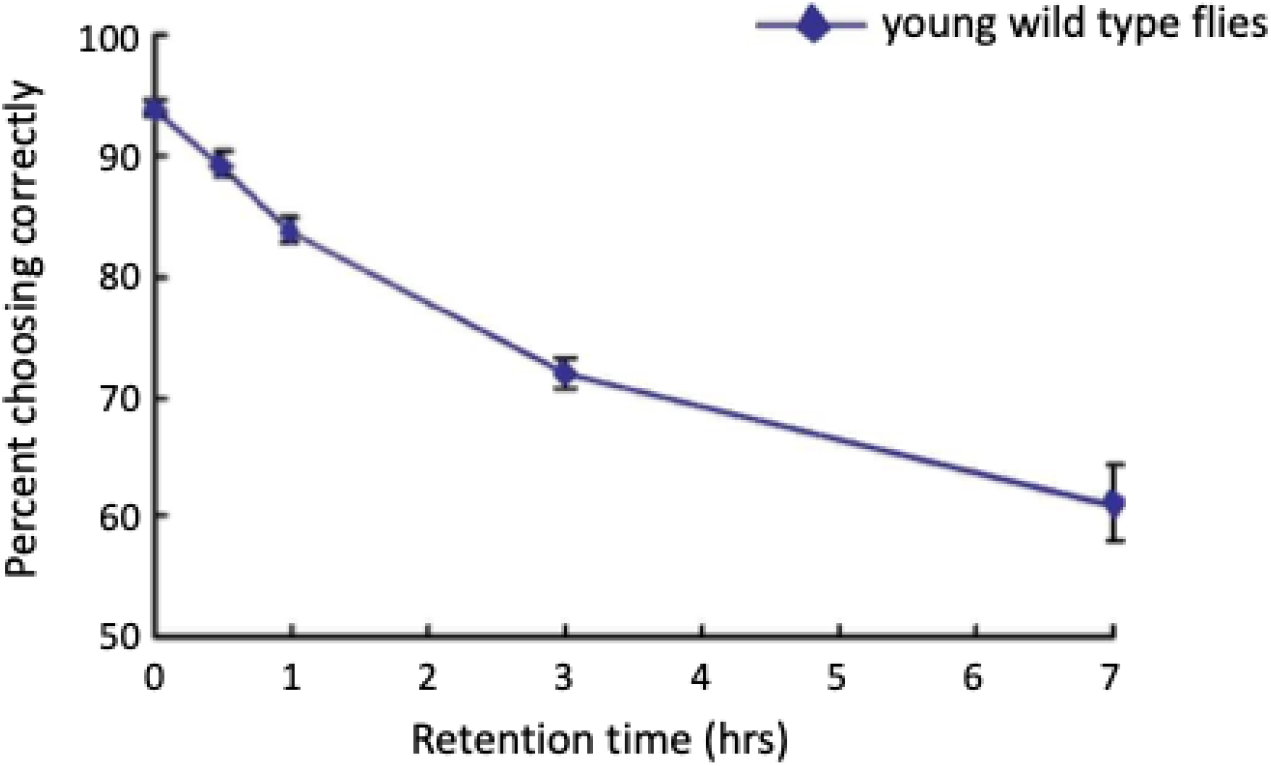
Memory retention curve. Flies were trained in an odor/shock association task as described in the text and tested at indicated times after training. R(C_1_), the percentage of flies choosing the correct odor, is plotted as a function of retention time. Data is used with permission from Tamura et al., 2003 (Tamura et al., 2003).

In *Drosophila*, learning and memory are often measured using an olfactory associative task in which flies are trained to associate an odor with pain (Quinn et al., 1974). A population of flies is exposed to an odor and at the same time exposed to aversive electrical shocks. Flies are next exposed to a second odor, this time in the absence of electrical shocks. Flies learn to associate the first, but not the second, odor with pain, and subsequently avoid this odor. Memory of this association can be tested by allowing trained flies to choose between the two odors in a T-maze (Tully & Quinn, 1985). Immediately after training, a large proportion of flies avoid the shock-paired odor and choose the unshocked odor. This proportion decreases as the time between training and testing increases, and this decrease is thought to reflect a time-dependent increase in forgetting. Thus, mutations that improve memory retention without increasing initial learning have been used to identify putative biochemical components regulating forgetting (Berry, Cervantes-Sandoval, Nicholas, & Davis, 2012; Berry & Davis, 2014; Davis & Zhong, 2017; Horiuchi, Yamazaki, Naganos, Aigaki, & Saitoe, 2008; Shuai et al., 2015; Shuai et al., 2010). In addition, *Drosophila* mutants with altered memory retention curves have been used to identify different memory phases, including initial learning (LRN), short-term memory (STM), middle-term memory (MTM), anesthesia-resistant memory (ARM), and long-term memory (LTM) (Tully & Quinn, 1985). These memory phases occur in a specific temporal order and persist for different durations. This temporal sequence suggests that forgetting rates may be related to the efficiency of transition between different memory phases. Thus, forgetting may consist of either a progressively smaller number of flies forming successive memory phases, or a progressive increase in uncertainty associated with successive memory phases.

In this study, we analyzed forgetting in *Drosophila* using behavioral analyses to determine whether it consists of a stochastic loss of memory in an increasing subset of flies, or whether it consists of a gradual reduction of memory strength that occurs throughout the population. Our results do not exclude the possibility that some flies spontaneously forget an association over time. However, our data indicate that there is a time-dependent decease in the probability that a fly with memory chooses the non-shocked (appropriate) odor. In other words, a significant aspect of forgetting in flies consists of a decrease in the ability of a memory to influence a behavior. We refer to this decrease as an increase in uncertainty.

## Results

In the *Drosophila* olfactory association training paradigm, odor concentrations are chosen such that naïve flies distribute 50:50 when given a choice between two odors. When flies are given this choice immediately after odor/shock training, approximately 95% of flies choose the non-shock paired odor (hereafter referred to as the correct odor), while approximately 5% choose the shock-paired odor (referred to as the incorrect odor). This result suggests two extreme possibilities. If a fly that learns the association always chooses the correct odor when tested immediately after training, 90% of flies must have learned the association while 10% did not. (The 95% of flies choosing the correct odor should consist of the 90% that learned and half of the 10% that didn’t.) On the other extreme, we consider the situation where all flies learn the association. In this case, learning must consist of a shift in the probability of choosing the correct odor from 50% to 95%. These two non-mutually exclusive possibilities can be generalized by the following mathematical model, which expresses behavior as a function of memory and memory strength:

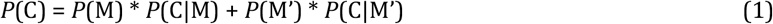

where *P*(C) is the probability that a fly chooses the correct odor, *P*(M) is the probability that a fly has memory of the odor association, *P*(C|M) is the probability that a fly that has memory chooses the correct odor, *P*(M’) is the probability that a fly has no memory of the association, and *P*(C|M’) is the probability that a fly that doesn’t have memory chooses the correct odor.

Flies that don’t form memories should distribute evenly between the odors, similar to naïve flies; thus, *P*(C|M’) = 0.5. In addition, flies should either have memory of the association or not; thus, *P*(M) + *P*(M’) = 1. Incorporating these constraints, equation (1) can be rewritten as:

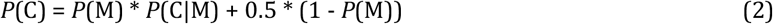

In the first situation described above, where a fly with memory always chooses the correct odor, *P*(C|M) = 1, and *P*(C) = (*P*(M) + 1)/2. In the second situation, where all flies learn the association, *P*(M) = 1 and *P*(C) = *P*(C|M).

We next considered what happens when forgetting occurs. Again, we first considered two extreme possibilities. If forgetting consists strictly of a stochastic loss of an associative memory in a subset of flies, *P*(M) should decrease over time, while *P*(C|M) will remain constant. On the other hand, if forgetting consists of a gradual reduction in certainty regarding an association that occurs in all flies that learned, *P*(C|M) will decrease while *P*(M) remains constant.

These two possibilities can be distinguished by testing flies for odor preferences at various times after training, and then separately retesting flies that chose the correct and incorrect odors in a second odor preference test immediately after the first test.

If forgetting consists of a stochastic loss of memory, trained flies should consist of two separable populations: one population that remembers and a second population that forgets. In this case, the initial odor preference test should separate flies into two non-equivalent populations. The population of flies that chose the correct odor should be enriched for flies that remember, while the population of flies that chose the incorrect odor should be highly enriched for flies that forget. When a second odor preference test is given immediately after the first, flies that chose correctly in the first test should again choose correctly in the second test at similar or higher probabilities. In contrast, flies that chose incorrectly in the first test should distribute between the odors at probabilities close to 50:50 in the second test. Thus, *P*(C_2_|C_1_) > *P*(C_2_|C_1_’), where *P*(C_2_|C_1_) is the probability that a fly that chose correctly in the first test chooses correctly in the retest, and *P*(C_2_|C_1_’) is the probability that a fly that chose incorrectly in the first test chooses correctly in the retest.

On the other hand, if forgetting consists strictly of a gradual reduction in certainty regarding the odor association that occurs in all or most flies in the population, *P*(M) should remain constant at a value close to 1, while *P*(C|M) should decrease over time. In this case, trained flies should not separate into two different populations. Instead, the probability of a fly choosing the correct odor should result from random or stochastic choices reflecting *P*(C|M) within a relatively homogeneous population. Thus, retesting flies that initially chose the correct odor and flies that initially chose the incorrect odor should yield odor preferences that are similar to each other, and to the results of the initial odor preference test. *P*(C_2_|C_1_) ≃ *P*(C_2_|C_1_’).

We first performed retest experiments on flies immediately after aversive olfactory training and observed that close to 90% of trained flies chose the non-shocked odor when tested immediately after training (Fig. 2). When we separately retested flies that initially chose correctly and flies that initially chose incorrectly, we found that both populations showed a significant preference for the non-shocked odor. This indicates that a significant percentage of flies that learned chose incorrectly in the original odor preference test. Thus, *P*(C|M) < 1.

**Figure 2.**
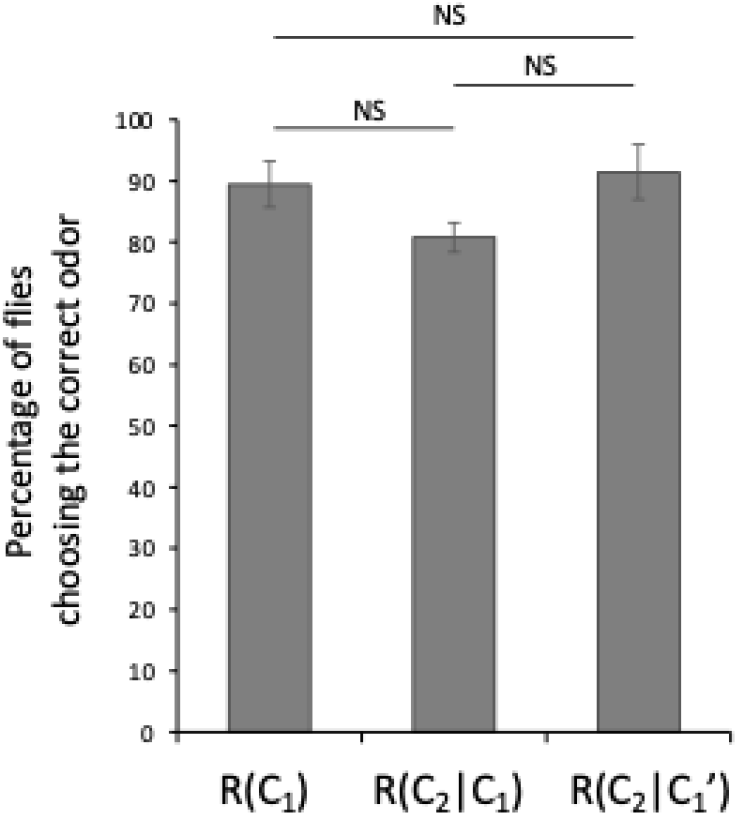
There is no significant difference in learning between flies that chose the correct or incorrect odors immediately after aversive olfactory conditioning. R(C_1_) refers to the percentage of flies that chose the correct (non-shocked) odor in the initial test immediately after training. R(C_2_|C_1_) refers to the percentage of flies that chose the correct odor in the retest after choosing the correct odor in the initial test, and R(C_2_|C_1_’) refers to the percentage of flies that chose the correct odor in the retest after initially choosing the incorrect odor. NS indicates p>0.5.

Next, to determine whether forgetting at 3 hrs consists of a decrease in the percentage of flies that remember, or a decrease in the probability that flies that remember choose the correct odor, we repeated the above experiment 3 hrs after training (Fig. 3). Unexpectedly, we found that a significantly higher percentage of flies that initially chose the incorrect odor chose the correct odor upon retest (compared to the percentage of flies that chose the correct odor in the initial test). Further, we found that a significantly lower percentage of flies that initially chose the correct odor chose the correct odor upon retest (compared to the percentage of flies that chose the correct odor in the initial test). It is extremely unlikely that flies that initially chose incorrectly have improved memory during the retest, suggesting that a non-associative effect must be occurring during the initial odor preference test that affects behavior during the retest.

**Figure 3.**
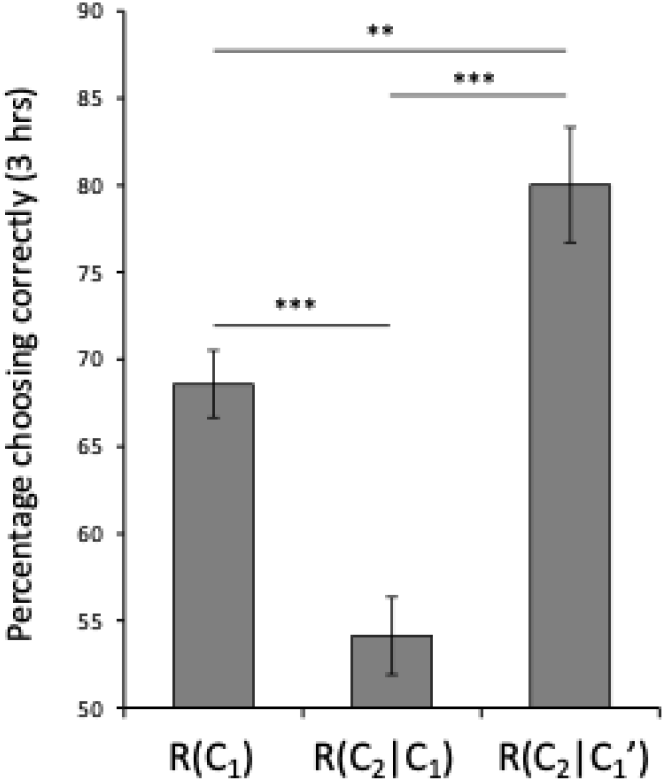
Non-associative effects influence odor retest assays 3 hrs after olfactory conditioning. The experiment is similar to that of Figure 1 except that the odor preference test and subsequent retest were performed 3 hrs after associative training. Flies that initially chose correctly, R(C_2_|C_1_), show significantly lower scores upon retest, while flies that initially chose incorrectly show significantly higher scores, R(C_2_|C_1_’), suggesting the existence of non-associative effects. **, p<0.01; ***, p<0.001.

To examine this possibility, we measured the behavior of naïve flies subjected to two consecutive odor preference tests (Fig. 4). As designed in our experimental paradigm, naïve flies distribute evenly between the two odors when initially tested for odor preference. However, upon retest, flies which initially chose one of the odors, octanol (Oct), preferred the second odor, methylcyclohexanol (MCH) upon retesting, while flies which initially chose MCH preferred Oct. Thus, flies tend to alternate their choice of odors, selecting odors that they had previously avoided when given a second chance. While we are not certain why flies have this tendency, we include several possible explanations for this behavior in the Discussion. Regardless of why flies behave in this curious manner, we refer to this behavior as an opposite preference tendency (T), and have included naïve controls in parallel with trained flies in all subsequent experiments.

**Figure 4.**
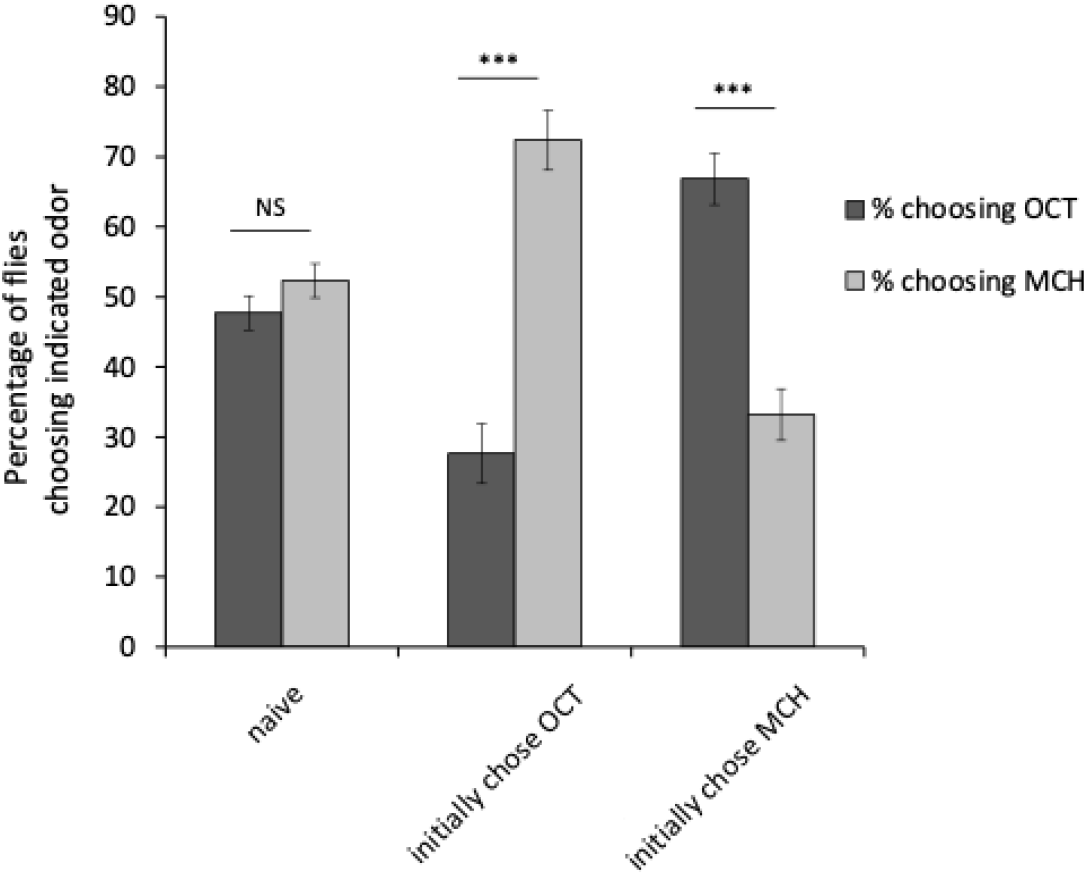
Opposite odor preference. While naïve flies distribute evenly between two odors (Oct and MCH) when initially tested for odor preference, those that initially chose Oct prefer MCH when retested, and those that initially chose MCH prefer Oct when retested. NS, p>0.05; **, p<0.01; ***, p<0.001.

We performed odor retest experiments for trained and naïve flies at various time points after training (Fig. 5). When we tested flies immediately after training (3 min time point), there were no significant differences in the percentage of flies choosing the correct odor during the first test or during retesting of flies that initially chose the correct or incorrect odors, and all of these scores were significantly different from the opposite preference tendency in naïve flies (Fig. 5A). This indicates that a proportion of flies that learned choose the incorrect odor during initial testing. Despite choosing incorrectly in the first test, these flies retain memory of the odor association and are able to choose the correct odor with a high probability on the second test.

**Figure 5.**
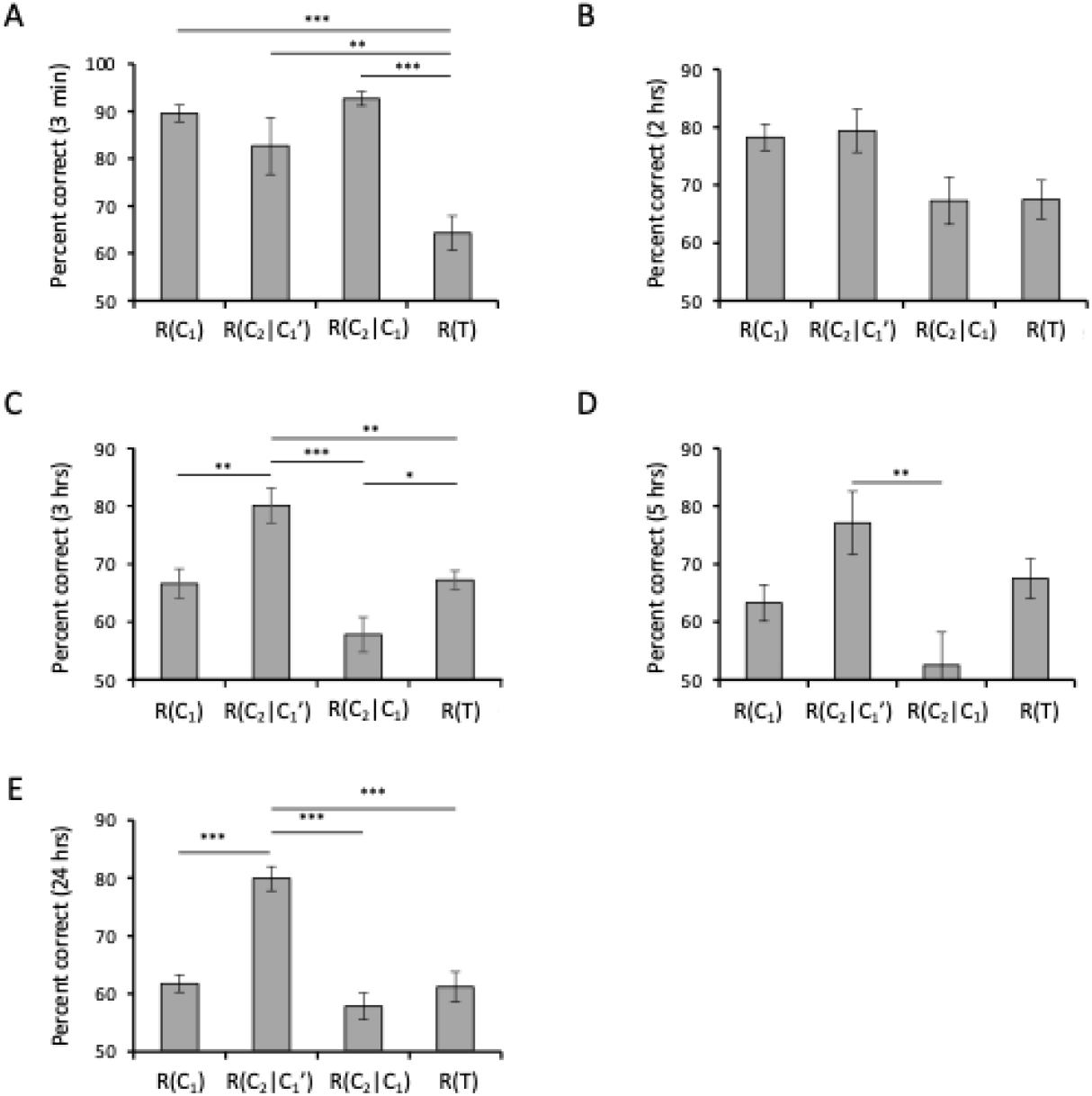
Initial odor preferences and retest odor preferences at 3 min (A), 2 hrs (B), 3 hrs (C), 5 hrs (D), and 24 hrs (E) after associative olfactory conditioning. The opposite odor preferences during retesting of untrained flies, R(T), is shown for comparison in each case. For A, B, C, and D, conditioning consisted of a single training trial, while for E, spaced conditioning was used. *, p<0.05; **, p<0.01; ***, p<0.001.

As the time interval between training and testing increases, the percentage of flies choosing the correct odor during the initial test decreases as flies gradually forget the association (Fig. 5A-D). Flies that chose correctly during this initial test show a decreased tendency to select the correct odor during the retest over time, while flies that chose the incorrect odor during the initial test show an increase in the probability of choosing the correct odor. It is important to keep in mind that for flies that initially chose the correct odor, the opposite preference tendency will work against the tendency for flies to choose the correct odor in the retest. For flies that initially chose the incorrect odor, the opposite preference tendency will enhance the tendency for flies to choose the correct odor in the retest. Thus, our data are consistent with a model in which odor preferences caused by training are strong immediately after training and dominate odor preference scores in both the initial tests and retests. As time after training increases, training-dependent associations decrease in strength, and retest scores tend to become the sum of memory-induced preferences and opposite odor preferences.

Flies can form long-term memories (LTMs), which can be measured 24 hrs after spaced training, a training protocol consisting of multiple (10x) training trials with 15 min rest intervals between each training. LTM has similarities with shorter forms of memory since it requires the same gross anatomical structures including the mushroom bodies and antennal lobes, but it is also distinct because it has different molecular requirements and uses different neuronal networks compared to short forms of memory. From our odor retest experiments, we find that LTM has qualitative similarities with short-lasting 3 hr and 5 hr memories (Fig. 5E). LTM increases the probability that flies will choose the correct odor.

However, flies with LTM will still choose the incorrect odor at some probability, despite retaining memory of the association and maintaining an increased probability of choosing the correct odor.

In order to examine how *P*(M) and *P*(C|M) are affected by forgetting, we used our measured values for R(C), R(C_2_|C_1_), R(C_2_|C_1_’), and R(T) as estimates of *P*(C), *P*(C_2_|C_1_), *P*(C_2_|C_1_’), and *P*(T) and calculated numerical ranges for *P*_*t*_(M), *P*_*t*_(C_1_|M), *P*_*t*_(C_2_|MC_1_), and *P*_*t*_(C_2_|MC_1_’) using equation (2) and the following equations.

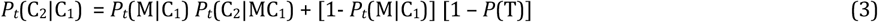

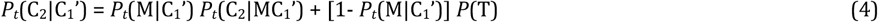

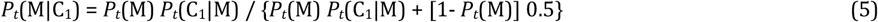

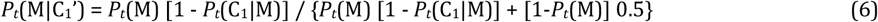

where,

*P*_*t*_(C_2_|C_1_) is the probability at time *t* that a fly will choose correctly in the second test given that it chose correctly in the initial test,

*P*_*t*_(M|C_1_) is the probability that a fly that chose correctly in the initial test has memory,

*P*_*t*_(C_2_|MC_1_) is the probability that a fly that has memory and chose correctly in the first test will choose correctly in the retest,

*P*(T) is the probability that a fly that chose an odor in the initial test will choose the opposite odor in the retest,

*P*_*t*_(C_2_|C_1_’) is the probability at time t that a fly will choose correctly in the second test given that it chose incorrectly in the first test,

*P*_*t*_(M|C_1_) is the probability that a fly that chose correctly in the initial test has memory, and *P*_*t*_(M|C_1_’) is the probability that a fly that chose incorrectly in the initial test has memory,

By restricting *P*_*t*_(M) to values between 0 and 1, and restricting *P*_*t*_(C_1_|M), *P*_*t*_(C_2_|MC_1_) and *P*_*t*_(C_2_|MC_1_’) to values between 0.5 and 1, we calculated possible ranges for *P*_*t*_(M), *P*_*t*_(C_1_|M), *P*_*t*_(C_2_|MC_1_) and *P*_*t*_(C_2_|MC_1_’). As seen in Figure 6A, the range of *P*(M) expands over time, encompassing both situations where *P*(M) remains constant and situations where *P*(M) decreases over time. In contrast, we find that values for *P*(C_1_|M) decrease over time (Fig. 6B). Overall, our data indicate that a major component of forgetting is a decrease in certainty, *P*(C|M), that occurs over time.

**Figure 6.**
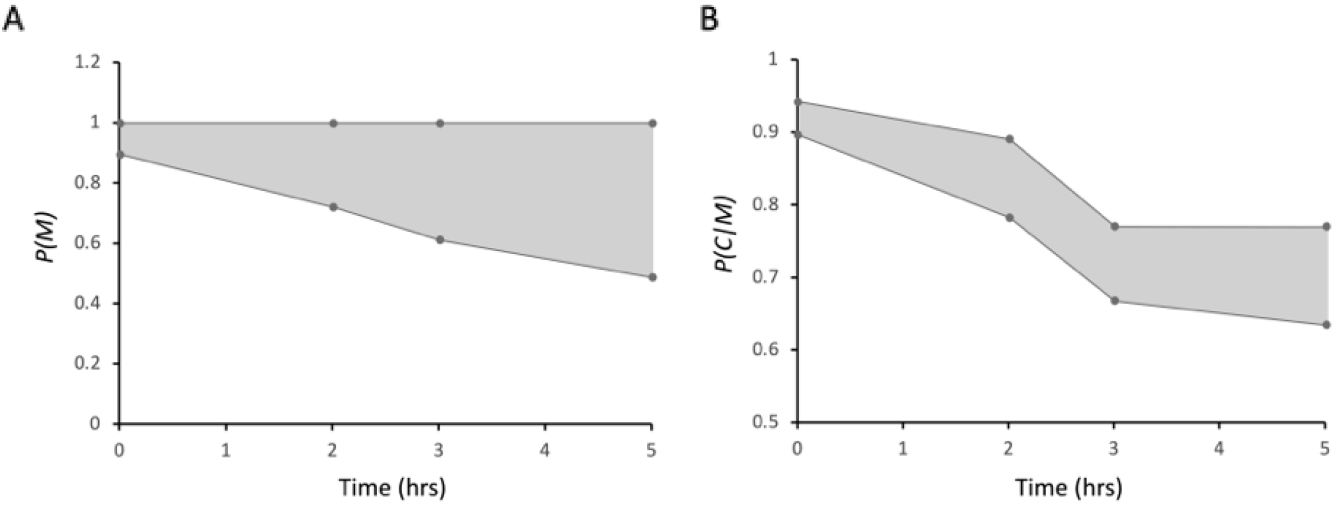
(A) Ranges for *P*_*t*_(M) at indicated times after training. (B) Ranges for *P*_*t*_(C|M) at indicated times after training.

Since aging in many species is associated with an increase in forgetting (Tamura et al., 2003), we next characterized short-term memory retention in old flies using the retesting procedure (Fig. 7). Overall, we observed reduced memory scores at early (3 min and 2 hrs) time points, and lesser, non-significant reductions at later (3 hr and 5 hr) timepoints in old flies compared to young flies. These reductions are consistent with age-dependent memory defects described in previous reports (Tamura et al., 2003; Yamazaki, Horiuchi, Miyashita, & Saitoe, 2010). We also observed reductions in retests for old flies that initially chose the correct odor compared to young flies (3 min and 2 hr time points). We did not observe significant differences in retests for old flies that initially chose the incorrect odor compared to young flies. Altogether, our results suggest that forgetting in old flies is qualitatively similar to forgetting in young flies, but with an accelerated rate of forgetting at early (0 hr and 2 hr) timepoints (Fig. 8).

**Figure 7.**
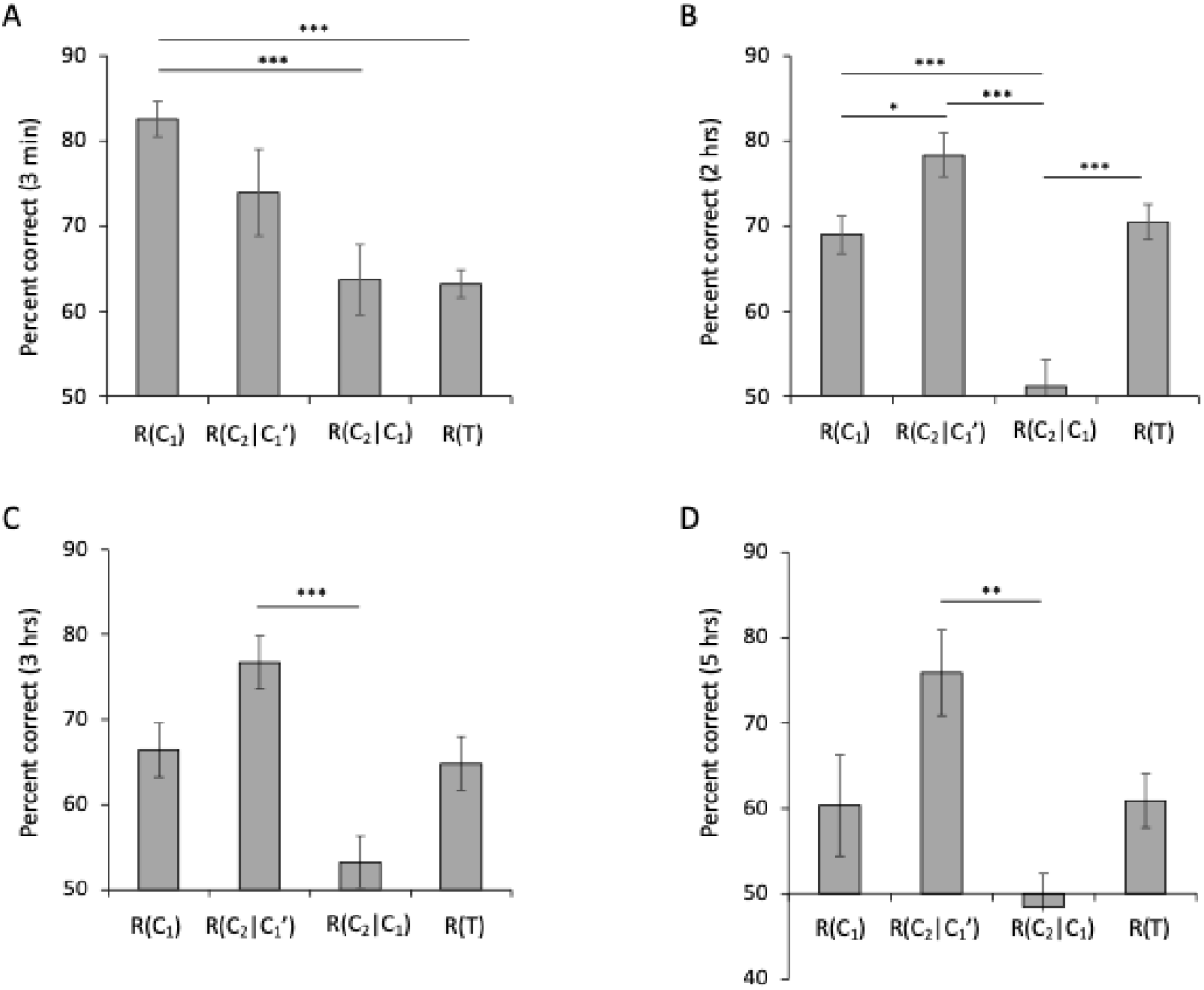
Initial odor preferences and retest odor preferences after olfactory conditioning of old (20-day-old) flies. *, p<0.05; **, p<0.01; ***, p<0.001.

**Figure 8.**
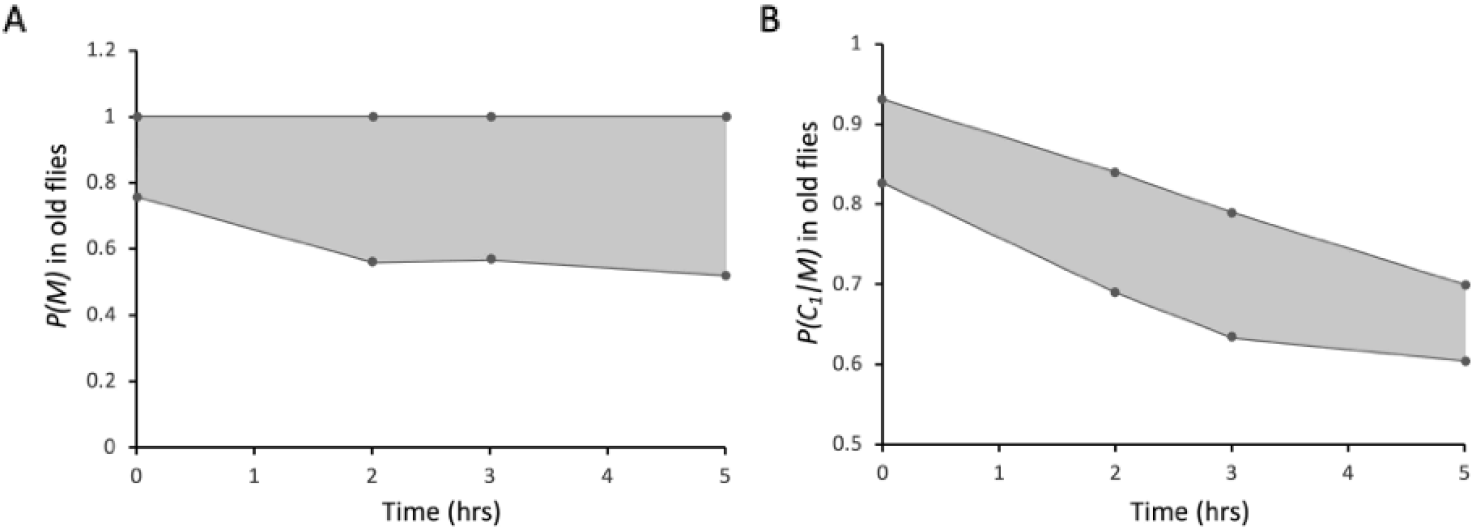
Ranges for (A) *P*_*t*_(M) and (B) *P*_*t*_(C|M) at indicated retention times for 20-day-old flies.

## Discussion

Although learning and memory have been studied extensively in *Drosophila*, it has been unclear precisely how memory decreases over time, and how forgetting occurs. For olfactory associative memories in *Drosophila*, flies form a simple association between a specific odor and either pain or rewards. Forgetting of this association could consist of a simple loss of memory or an increase in uncertainty regarding this association. Our study demonstrates that a large component of forgetting consists of a general decrease in the probability that a fly that remembers will choose the correct odor. This can be thought of as a time-dependent decrease in memory strength or an increase in uncertainty.

In *Drosophila*, a general memory retention curve consists of the summation of various different memory phases, including LRN, STM, MTM, ARM, and LTM (Tully et al., 1990; Tully, Preat, Boynton, & Del Vecchio, 1994). Our data suggest that forgetting does not consist strictly of a decrease in the probability that one memory phase is converted into a subsequent memory phase. In other words, it is unlikely that the main component of forgetting consists of subsequently smaller subset of flies forming STM, MTM, ARM, and LTM after learning. Instead, our data suggest that later memory phases differ from earlier memory phases in their ability to motivate flies to choose the correct odor.

Previous studies have tried to use behavioral tests to separate flies that learned from flies that did not (Mery, 2007). However, our work indicates that this strategy may have difficulties because we do not find significant differences in memory between flies that choose the correct odor and those that choose the incorrect odor. Our results are somewhat surprising since testing is almost exclusively used in human education and in mammalian learning studies to evaluate learning and memory in individuals. How are memory and forgetting in our experiments different from memory and forgetting in mammals? Flies used in our experiments are from a homogeneous genetic background and were raised in identical conditions. More importantly, our olfactory associative training paradigm is not an operant conditioning paradigm. Flies do not have a choice during training and are forcibly exposed to odors for one minute each and shocked 12 times during exposure to the shock-paired odor. Thus, factors such as motivation to learn and amount of study are normalized in our study. In this situation, our results suggest that forgetting occurs similarly in all individuals in a population. When training is increased (compare results from spaced training (figure 4E) versus single cycle (figure 4C,D)), flies retain memory for greater time periods, such that memory 24 hrs after 10x spaced training is similar to 3 or 5 hr memory after single cycle training. This indicates that in flies as well as humans, the amount of training or study strongly affects the rate of forgetting.

During our studies, we observed that naïve flies distribute evenly when given a choice between the two odors used in our study. However, when flies were retested with the two odors after an initial odor choice, they preferentially chose the opposite odor to the one they had initially chosen. This behavior may have some similarities to a behavior known as Buridan’s paradigm in which a fly will continuously walk back and forth between two dark lines at opposite ends of an otherwise featureless arena (Colomb, Reiter, Blaszkiewicz, Wessnitzer, & Brembs, 2012; Han, Wei, Tseng, & Lo, 2021). Buridan’s paradigm suggests that flies tend to alternate when given two choices instead of focusing on one or the other. Alternatively, it is possible that the T-maze used during testing is slightly aversive to flies. Thus, when flies choose one odor in one arm of the maze, this odor becomes associated with an aversive experience, which then affects the retest. Consistent with this idea, we find that the opposite preference behavior becomes slightly weaker when the plastic arms of the T-maze are coated with filter paper, and is abolished when the arms of the T-maze are coated with a sucrose reward (data not shown).

Why does forgetting consist of a decrease in the probability that a memory induces a behavior? It seems likely that learning and memory originally evolved to increase the survival of an organism. If this is the case, forgetting should also have evolved as a calculated survival response to a risk. In contrast to the survival strategy of humans, who have relatively few children per parent and raise each child carefully, the survival strategy of flies involves mass production of progeny. In this situation, having a certain population of progeny that chooses the incorrect odor may be evolutionarily beneficial, since it can prevent catastrophic loss of the entire population in cases where a situation rapidly changes. Thus, the increase in the proportion of flies choosing the incorrect odor may reflect a calculation that the reliability of a memory as a future predictor decreases as a memory gets older. Preserving the actual memory (M) itself may be important for future events, such as accelerated reinstatement of a behavior if the odor-shock association is re-experienced (sensitization). Thus, decreasing *P*(C|M) over time may be an optimal method of maintaining some memory of an association while at the same time decreasing its effect on behavior as time progresses. It is fascinating to consider that this type of survival strategy may have evolved into more complex forgetting in humans and other mammals.

Memory retention curves in flies resemble those of humans, first described by Ebbinghaus in 1885 (Ebbinghaus, 1885; Ebbinghaus, 2013). While the time scales are vastly different between these curves, their overall shapes are similar, with a rapid reduction in memory at early time points that becomes more gradual over time. This suggests that forgetting in flies shares aspects of forgetting with humans. Consistent with this idea, aging affects memory retention in flies, similar to humans and other animals, again suggesting similarities between forgetting in flies and humans.

## Materials and Methods

### Fly lines and maintenance

*w*(CS) flies were used in all experiments in this study. Flies were raised at 25 °C and 60% humidity on at 12 hr: 12 hr light-dark cycle. 3 to 5-day-old flies were used for young flies, and 20-day-old flies were used for old flies. Flies were aged in food vials containing ~50 flies each and were transferred into new vials every two to three days.

### Olfactory associative training protocols

The aversive olfactory associative training procedure has been previously described (Tamura et al., 2003). Briefly, ~100 flies were placed in a training chamber where they were exposed to an odor (the shock-paired conditioned stimulus, or CS+) for 1 min and simultaneously shocked 12 times for 1.5 secs at 60 volts every 5 secs. After a 45 sec air purge, flies were exposed to a second odor (the unpaired odor or CS-) for 1 min in the absence of electrical shocks. If flies learned, they should associate the CS+, but not the CS-, with pain. Spaced training consisted of ten repeated trainings with 15 min rest intervals between each training cycle. All behavioral experiments were repeated 8 to 16 times for each time point, alternating 1-octanol and 4 methyl cyclohexanol for the CS+ and CS-odors.

### Testing protocols

For testing, flies were placed in an elevator and manually moved to the choice point of a T-maze where they were allowed to choose between the CS+ and CS-odors for 90 seconds. Immediately after the initial test, flies that chose each odor were retested with the same odors.

### Mathematical modelling

We investigated time-related changes in memory [*P*_*t*_(M)] and certainty about odor associations [*P*_*t*_(C_1_|M)] based on equations (2-6) described in the results. We used measured ratios, R_t_(C1), R_t_(C2|C1’), R_t_(C2|C1), and R(T), as estimates for *P*_*t*_(C1), *P*_*t*_(C2|C1’), *P*_*t*_(C2|C1), and *P*(T), leaving *P*_*t*_(M), *P*_*t*_(C_1_|M), *P*_*t*_(C_2_|MC_1_), and *P*_*t*_(C_2_|MC_1_’) as unknowns.. Since the number of variables exceeded the number of equations, we were unable to obtain precise values, but by restricting *P*_*t*_(M) to values between 0 and 1, and restricting *P*_*t*_(C_1_|M), *P*_*t*_(C_2_|MC_1_), and *P*_*t*_(C_2_|MC_1_’) to values between 0.5 and 1, we calculated possible ranges for *P*_*t*_(M), *P*_*t*_(C_1_|M), *P*_*t*_(C_2_|MC_1_) and *P*_*t*_(C_2_|MC_1_’) that were used to generate figures 6 and 8. All modeling and calculations were performed using Microsoft Excel and GraphPad Prism software.

## Acknowledgments

This work was supported by Japan Society for Promotion of Science grants to J.H. (21K06403,18K06496) and M.S. (19H01013, 21K18238).

